# HSF2 protects against proteotoxicity by maintaining cell-cell adhesion

**DOI:** 10.1101/506881

**Authors:** Jenny Joutsen, Alejandro J. Da Silva, Marek A. Budzynski, Jens C. Luoto, Aurelie de Thonel, Jean-Paul Concordet, Valérie Mezger, Délara Sabéran-Djoneidi, Eva Henriksson, Lea Sistonen

## Abstract

Cellular ability to maintain proper protein homeostasis (proteostasis) is essential for survival upon protein-damaging conditions. Heat shock transcription factor 2 (HSF2) is one of the human HSFs activated in response to proteotoxic stress. HSF2 is dispensable for cell survival during acute heat stress, but its amount and DNA-binding activity increase under prolonged proteotoxic stress conditions, such as proteasome inhibition. Nevertheless, the specific role(s) of HSF2 and the global HSF2-dependent gene expression profile during sustained stress have remained elusive. We found that HSF2 is required for cell survival during prolonged proteotoxicity, as shown by treating wild-type and HSF2-deficient human osteosarcoma U2OS cells with the proteasome inhibitor Bortezomib. Strikingly, our RNA-seq analyses revealed that HSF2 disruption leads to marked downregulation of cadherin superfamily genes and subsequent functional impairment of cadherin-mediated cell-cell adhesion. We propose HSF2 as a key regulator of genes belonging to the cadherin superfamily. We also demonstrate that HSF2-dependent downregulation of cadherin-mediated cell-cell adhesion predisposes U2OS cells to Bortezomib-induced proteotoxic stress. In conclusion, we show that by maintaining cell-cell adhesion HSF2 is essential for cell survival and thereby we describe a novel regime in the HSF-mediated protection against stress-induced protein damage.

## INTRODUCTION

The cells in a human body are constantly exposed to environmental stressors, which challenge the maintenance of protein homeostasis (proteostasis). To survive insults that disturb proteostasis, cells rely on a selection of protective mechanisms that can be launched upon stress exposures. The heat shock response is a well-conserved stress protective pathway that is induced in response to cytosolic protein damage and mediated by heat shock transcription factors, HSFs [1]. Upon activation, HSFs oligomerize, accumulate in the nucleus, and bind to their target heat shock elements (HSEs) at multiple genomic loci [2–4]. The canonical HSF target genes encode molecular chaperones, such as heat shock proteins (HSPs), which assist in the maintenance of correct protein folding environment by refolding the misfolded proteins or directing them to protein degradation machineries [5]. In addition to regulating the transcriptional response to stress, HSFs are important in a variety of other physiological and pathological processes and the repertoire of HSF target genes extends greatly beyond the HSPs [6–10].

The human genome encodes six HSF family members (HSF1, HSF2, HSF4, HSF5, HSFX and HSFY), of which HSF1 and HSF2 are the most extensively studied [13]. Although HSF1 and HSF2 are homologous in their DNA-binding domains, they share only few similarities in the tissue expression patterns, regulatory mechanisms, and signals that stimulate their activity [1, 14]. HSF2 is an unstable protein and its expression is highly context-dependent, fluctuating in different cell and tissue types [15, 16], developmental stages [17, 18], and during the cell cycle [19]. Consequently, regulation of HSF2 protein levels has been generally accepted as the main determinant of its DNA-binding capacity [20, 21]. In mouse, lack of HSF2 results in developmental defects particularly associated with corticogenesis and spermatogenesis [22, 23]. HSF1 is essential for HSP expression and cell survival in a variety of acute stress conditions [13], whereas HSF2 is downregulated in cells exposed to acute heat stress [24] and has only a modulatory role in the regulation of HSP expression [4, 25, 26]. Interestingly, HSF2 is upregulated and acquires DNA-binding activity in cells exposed to proteasome inhibition induced by lactacystin or MG132 [20, 27, 28].

The ubiquitin-proteasome system is one of the main cellular mechanisms regulating protein turnover, thereby affecting multiple aspects of cell physiology, such as signal transduction and apoptosis [29, 30]. Due to the fundamental function in cell physiology, the proteasome complex has emerged as an important target for anti-cancer therapy [31]. The most common drug to inhibit proteasome function is Bortezomib (PS-341, V?LCADE^®^), which is currently used as a standard treatment in hematological malignancies [32]. Bortezomib is a dipeptide boronic acid derivative that targets the chymotrypsin-like activity of the 26S proteasome, causing progressive accumulation of damaged proteins [32–34]. By exposing human blood-derived primary cells to clinically relevant concentrations of Bortezomib, Rossi and colleagues demonstrated that prolonged proteasome inhibition results in a remarkable upregulation of HSF2 at both mRNA and protein levels [35]. That study also showed that HSF2, together with HSF1, localizes to the promoters of *HSP70* and *AIRAP* (zinc finger AN1-type domain 2a), but that HSF2 is not required for the induction of these genes upon Bortezomib treatment [35, 36]. Interestingly, proteasome inhibition has been shown to be severely toxic to HSF2-deficient mouse embryonic fibroblasts [37], indicating that HSF2 is required for protection against proteotoxicity although its distinct role and the global HSF2-dependent gene expression program have remained unknown.

In this study, by comparing Bortezomib-treated wild-type and HSF2 knock-out human osteosarcoma U2OS cells, we show that HSF2 is essential for cell survival upon prolonged Bortezomib-induced proteotoxicity. To our surprise, the RNA-seq analyses revealed that HSF2 disruption results in a profound downregulation of genes belonging to the cadherin superfamily and functional impairment of the cadherin-mediated cell-cell adhesion. Furthermore, we show that inhibition of cadherin-mediated cell-cell adhesion predisposes human U2OS cells to Bortezomib-induced cell death. Taken together, our results demonstrate that by maintaining cadherin-mediated cell-cell adhesion, HSF2 acts as an important pro-survival factor during sustained proteotoxic stress.

## RESULTS

### U2OS cells lacking HSF2 are predisposed to Bortezomib-induced proteotoxicity

To explore the role of HSF2 during prolonged proteotoxic stress, we first examined the expression and cellular localization of HSF2 during Bortezomib (BTZ) treatment. Human osteosarcoma U2OS WT cells were treated with different concentrations of BTZ (0-100 nM) for 6 or 22 h and HSF2 protein levels were examined with immunoblotting. The time points were selected to assess both the short-term and the long-term exposure to BTZ. Interestingly, HSF2 was slightly upregulated already at the 6-h time point, whereas at 22 h its expression was highly elevated (Fig. 1A). Indirect immunofluorescence of U2OS WT cells revealed that HSF2, which is known to be both cytoplasmic and nuclear [38, 39], resides in the nucleus already under control conditions and the nuclear localization was further enhanced during BTZ treatment (Fig. 1B). These results are in line with a previous report using human blood-derived primary cells [35] and show that also malignant U2OS cells respond to BTZ treatment with marked increase in HSF2 levels and accumulation in the nucleus.

**Figure 1.**
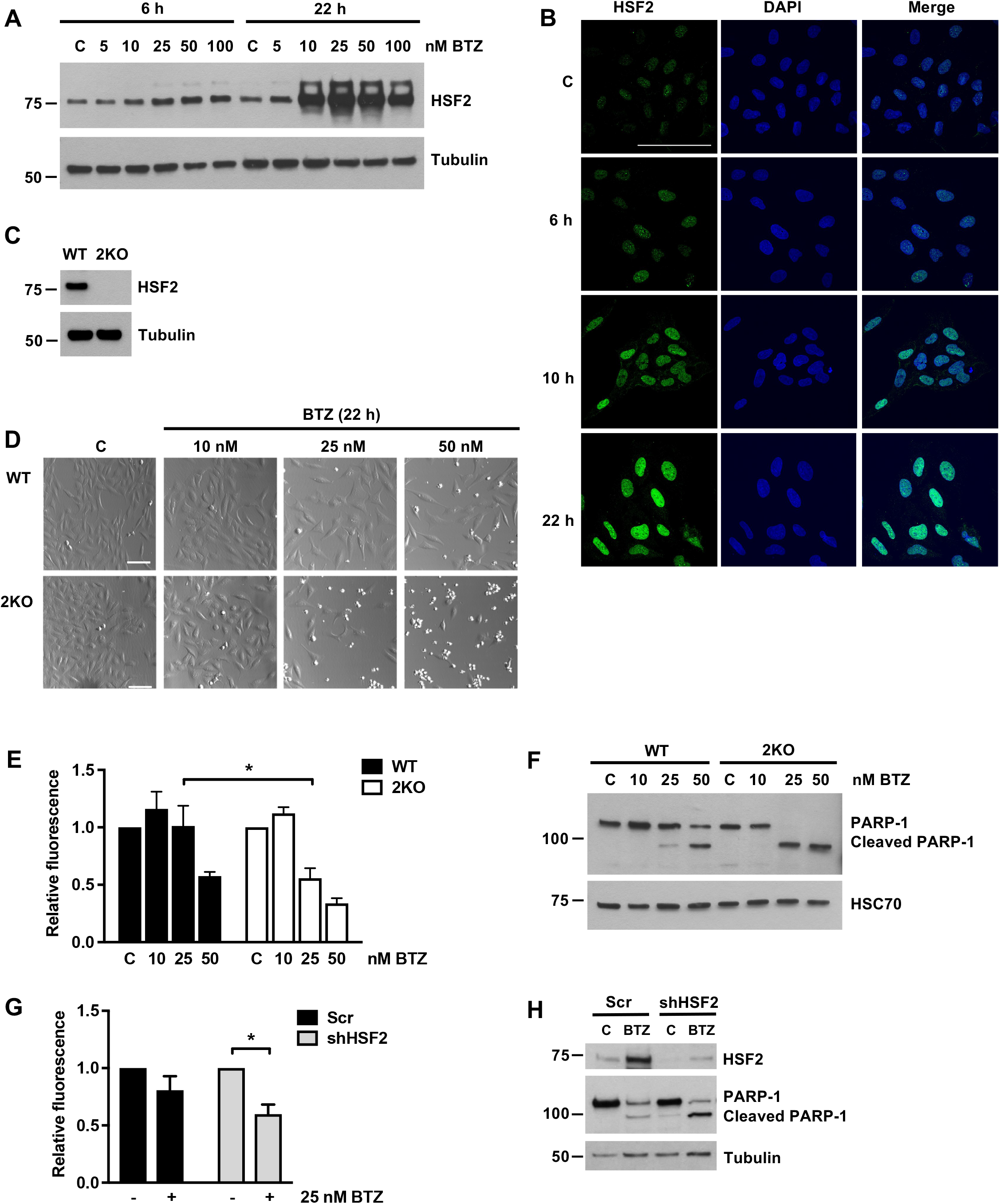
HSF2 is required for cell survival upon prolonged Bortezomib (BTZ) treatment. **(A)** Immunoblot analysis of HSF2 expression. U2OS WT cells were treated with indicated concentrations of BTZ for 6 or 22 h. Control cells were treated with DMSO. Tubulin was used as a loading control. **(B)** Confocal microscopy images of HSF2 immunofluorescence staining. U2OS WT cells were plated on cover slips and treated with 25 nM BTZ for 6, 10, or 22 h. Control cells were treated with DMSO. Samples were fixed and stained with anti-HSF2 antibody. DAPI was used for DNA detection. The overlay of HSF2 and DAPI maximum intensity projection signals is shown in merge. Scale bar 100 μm. **(C)** Immunoblot analysis of HSF2 expression in U2OS WT and HSF2 KO (2KO) cells. Tubulin was used as a loading control. **(D)** Bright-field microscopy images of WT and 2KO cells treated with indicated concentrations of BTZ for 22 h. Control cells were treated with DMSO. Scale bar 100 μm. **(E)** Calcein AM assay of WT and 2KO cells treated as in D. Relative fluorescence was calculated against each respective control that was set to 1. The data is presented as mean values of at least three independent experiments + SEM, *p < 0.05. **(F)** Immunoblot analysis of PARP-1. Cells were treated as in D. HSC70 was used as a loading control. **(G)** Calcein AM assay of U2OS WT cells transfected with Scr or HSF2-targeting shRNA constructs [25] and treated with 25 nM BTZ for 22 h. Relative fluorescence was calculated against each respective control that was set to 1. The data is presented as mean values of three independent experiments + SEM, *p < 0.05. **(H)** Immunoblot analysis of HSF2 and PARP-1. Cells were transfected and treated as in G. Tubulin was used as a loading control.

Due to the prominent upregulation of HSF2 in cells treated with BTZ, we next asked if HSF2 is required for the cellular survival under these stress conditions. We generated a U2OS *HSF2* knock-out cell line (2KO hereafter), where HSF2 expression was abolished by mutating the first exon of the *HSF2* gene using the CRISPR-Cas9 method. In these cells the protein expression of HSF2 was completely abrogated (Fig. 1C). U2OS WT and 2KO cells were treated with indicated concentrations of BTZ for 22 h and examined with microscopy. We observed a dramatic difference in the viability of the WT and 2KO cells, since the cells lacking HSF2 exhibited an apoptotic non-adherent phenotype in concentrations where WT cells remained adherent (Fig. 1D). Quantitative cell viability measurements confirmed that the survival of 2KO cells was significantly impaired upon BTZ treatment (Fig. 1E). Furthermore, 2KO cells accumulated more cleaved PARP-1 than WT cells, demonstrating a more pronounced activation of apoptosis [40] (Fig. 1F). Similar results were obtained with another *HSF2* knock-out cell line (2KO#2 hereafter) (Fig. S1A), confirming that the observations are not specific for a single clone. In addition to BTZ, treatments with MG132, a well-established proteasome inhibitor, and amino acid analogue L-canavanine, which causes protein misfolding when incorporated into nascent peptide chains, clearly reduced viability of HSF2-deficient cells (Fig. S1B-F). Previously, *HSF2*-null mouse embryonic fibroblasts (*Hsf2^-/-^* MEFs) have been shown to be sensitized to MG132 treatment as well as to sustained febrile-range thermal stress [37, 41]. Altogether these results indicate that HSF2 is an important survival factor in both murine and human cells when exposed to prolonged proteotoxic stress.

To verify that the decreased survival of 2KO cells was not caused by off-target effects of the CRISPR-Cas9 method, we transfected the U2OS WT cells with scramble (Scr) or HSF2-targeting shRNA plasmids and treated the cells with BTZ for 22 h. In accordance with the results obtained with 2KO cells, transient HSF2 downregulation significantly reduced cell viability upon BTZ treatment (Fig. 1G). Moreover, HSF2 downregulation enhanced the progression of apoptosis, which was detected by increased accumulation of cleaved PARP-1 (Fig. 1H). Hence, we conclude that HSF2 is required for cell survival upon BTZ-induced proteasome inhibition.

In contrast to HSF2, which has been found to be downregulated in a subset of human cancers [12], both the expression and activity of the closely related HSF1 are increased in a majority of studied cancer types [1]. A marker for HSF1 activation is phosphorylation of serine 326 (pS326) [7, 42], and HSF1 expression and pS326 were recently established as requirements for multiple myeloma cell survival during BTZ treatment [36]. Therefore, we examined whether the decreased survival of 2KO cells was due to impaired HSF1 expression or phosphorylation upon proteasome inhibition. U2OS WT and 2KO cells were treated with BTZ or MG132 and the HSF1 protein levels and S326 phosphorylation status were analyzed with immunoblotting. Interestingly, no difference in HSF1 expression or S326 phosphorylation between WT and 2KO cells was observed upon proteasome inhibition (Fig. S1G). These results demonstrate that although HSF1 is an essential survival factor during acute stress [1], it is not sufficient to protect cells during prolonged proteotoxicity.

### Bortezomib-induced heat shock response is not compromised in HSF2-deficient cells

Similarly to many other surveillance transcription factors, such as p53 [43], HIF-1α [44] and Nrf2 [45], HSF2 is an unstable protein under normal growth conditions [24]. HSF2 expression has been shown to fluctuate in response to stress exposure, tumor progression, and during the cell cycle [12, 19, 24], and high expression levels of HSF2 correlate with its increased DNA-binding activity [20, 47]. Due to the massive increase in nuclear HSF2 levels upon BTZ treatment (Fig. 1B), we investigated if the impaired survival of 2KO cells is caused by misregulation of HSF2 target genes. U2OS WT and 2KO cells were treated with 25 nM BTZ for 6 or 10 h (Fig. 2A) and the global gene expression profiles were analyzed with RNA-seq. The selected time points represent sub-lethal proteotoxic stress conditions, at which the cell viability is not yet compromised (Fig. S2A). Before mRNA purification, the knock-out phenotype was confirmed with immunoblotting (Fig. S2B). Notably, stress-inducible hyperphosphorylation of HSF1 [47] and increased HSP70 expression were observed in both WT and 2KO cells (Fig. S2B). To identify the HSF2-dependent target genes, we first compared the inducible gene expression profiles between WT and 2KO cells in response to 6 h and 10 h BTZ treatment (Fig. 2A). Differentially expressed (DE) genes were determined with the Bioconductor R package Limma [48], with fold change ≥ 3 and false discovery rate (FDR) < 0.001, from quadruplet samples that correlated well to each other (Fig. S2C). According to the analysis, BTZ treatment resulted in a significant upregulation and downregulation of genes in WT (> 600 and > 300, respectively) and 2KO (> 500 and > 200, respectively) cells (Fig. 2B, Table S1). The complete dataset is available at Gene Expression Omnibus under accession number: GSE115973.

**Figure 2.**
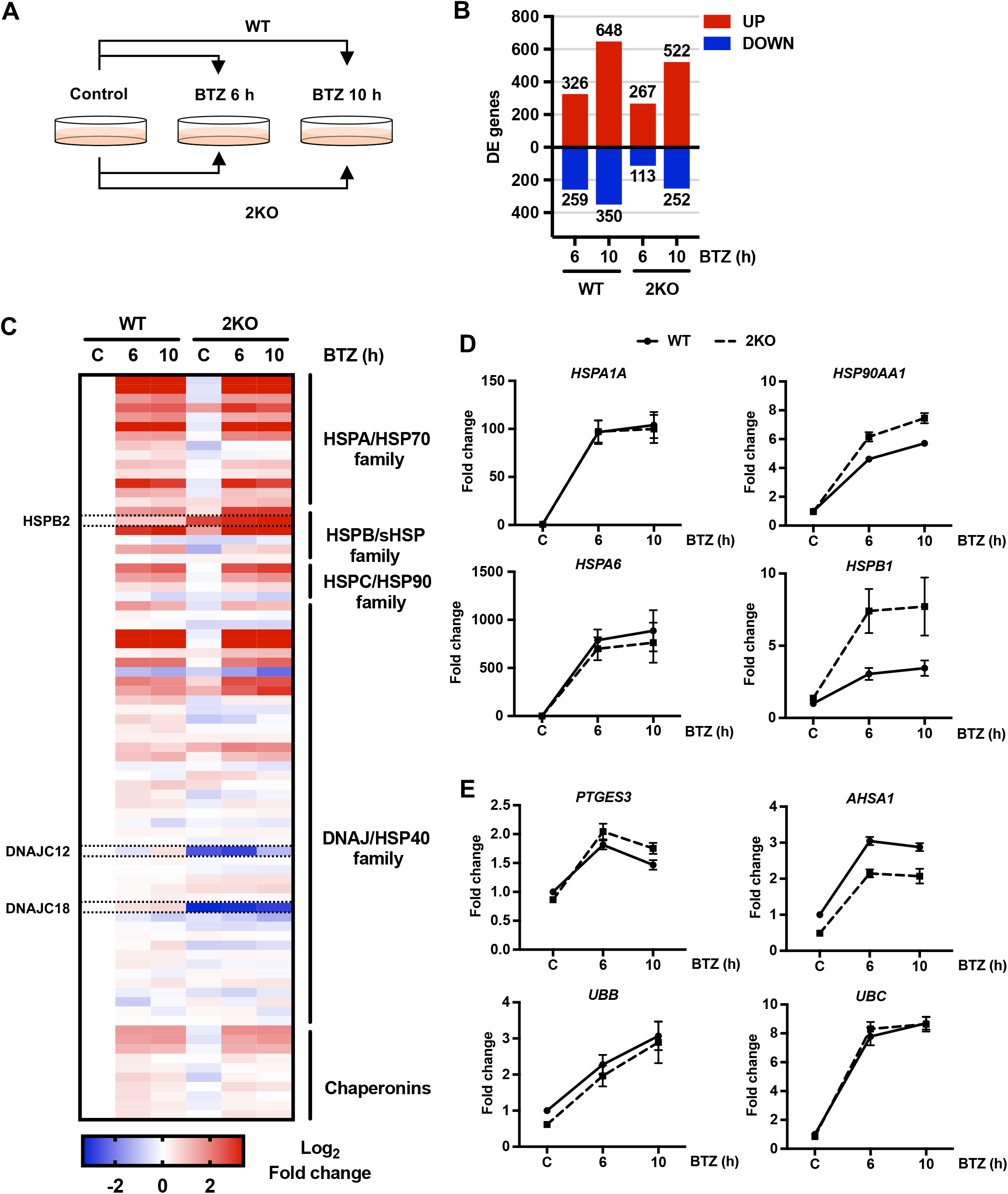
HSF2 is not required for the heat shock response during BTZ-induced proteotoxic stress. **(A)** A schematic overview of the RNA-seq experiment outline. U2OS WT and HSF2 KO (2KO) cells were treated with 25 nM BTZ for 6 or 10 h. Control cells were treated with DMSO. After treatments, mRNA was extracted and analyzed by RNA-seq. Experiments were performed in biological quadruplets. The arrows depict comparison pairs. **(B)** Differentially expressed (DE) genes in each comparison pair were determined with Bioconductor R package Limma [48] (FC ≥ 3, FDR < 0.001). The upregulated and downregulated genes in a given comparison pair are indicated with red and blue bars, respectively. **(C)** Normalized gene expression data for human heat shock proteins, as defined in [49], was used to calculate the fold change of each gene in relation WT control sample. The data is presented as a heatmap of log2 transformed values and it was generated with GraphPad Prism7. Examples of genes that exhibit a divergent expression pattern are framed. (D, E) mRNA expression levels of selected heat shock proteins (*HSPA1A, HSP90AA1, HSPA6*, and *HSPB1*) **(D)**, HSP90 co-chaperones (*PTGES3* and *AHSA1*), and stress-responsive ubiquitin genes (*UBB* and *UBC*) **(E)** determined with RNA-seq. The data is presented as mean values ± SEM relative to WT control sample that was set to 1.

The HSF-regulated heat shock response is one of the main cellular survival pathways induced by proteotoxic stress and it is characterized by simultaneous upregulation of multiple genes essential for maintaining the correct protein folding environment [1, 4]. To examine whether the heat shock response is compromised in 2KO cells, we analyzed the inducible expression patterns of all human molecular chaperone genes [49], in WT and 2KO cells treated with BTZ. Intriguingly, the chaperone expression profiles of WT and 2KO cells were nearly identical, and only *HSPB2, DNAJC1,2* and *DNAJC18* exhibited clearly distinct expression patterns in 2KO cells (Fig. 2C). A closer examination of the RNA-seq data for the expression of selected chaperone genes, i.e. *HSPA1A* (HSP70), *HSP90AA1* (HSP90), *HSPA6* (HSP70B’), and *HSPB1* (HSP27), revealed equal or even higher expression levels in 2KO cells than in WT cells (Fig. 2D). HSF2 has been previously shown to negatively modulate HSF1-mediated activation of specific HSP targets [19, 25, 26], which likely explains the higher expression of *HSPB1* and *HSP90AA1* in 2KO cells upon BTZ treatment (Fig. 2D). In response to proteotoxic stress, HSF2 localizes also to the promoters of HSP90 co-chaperone and polyubiquitin genes [2]. To study whether the regulation of these genes was disturbed in 2KO cells, HSP90 co-chaperones *PTGES3* (p23) and *AHSA1* (AHA1) as well as polyubiquitin genes (*UBB* and *UBC*) were examined from our RNA-seq data. Since no significant differences were observed in the expression patterns of any of these genes (Fig. 2E), we conclude that lack of HSF2 does not compromise the induction of the heat shock response upon BTZ treatment.

### Disruption of HSF2 leads to misregulation of cell adhesion associated genes

To determine the differences between the WT and 2KO cells, we examined the DE genes in 2KO : WT comparison pair at each experimental time point (control, 6, and 10 h) (Fig. 3A). Despite the stringent cutoff criteria (FC ≥ 3, FDR 0.001), 2KO cells exhibited significant misregulation of 819 genes already under normal growth conditions (2KO C : WT C), and the proportion of upregulated and downregulated genes remained similar throughout the BTZ treatments (2KO 6 h : WT 6 h, 2KO 10 h : 2KO 10 h) (Fig. 3B, Table S1). GO term analysis of the misregulated genes revealed a specific enrichment of terms related to cell adhesion and cell-cell adhesion *via* plasma membrane adhesion molecules (Fig. 3C, Table S1). Similar GO terms among the comparison pairs implied that the genes misregulated in 2KO cells are tightly linked to cellular adhesion properties both under control and stress conditions (Fig. 3C).

**Figure 3.**
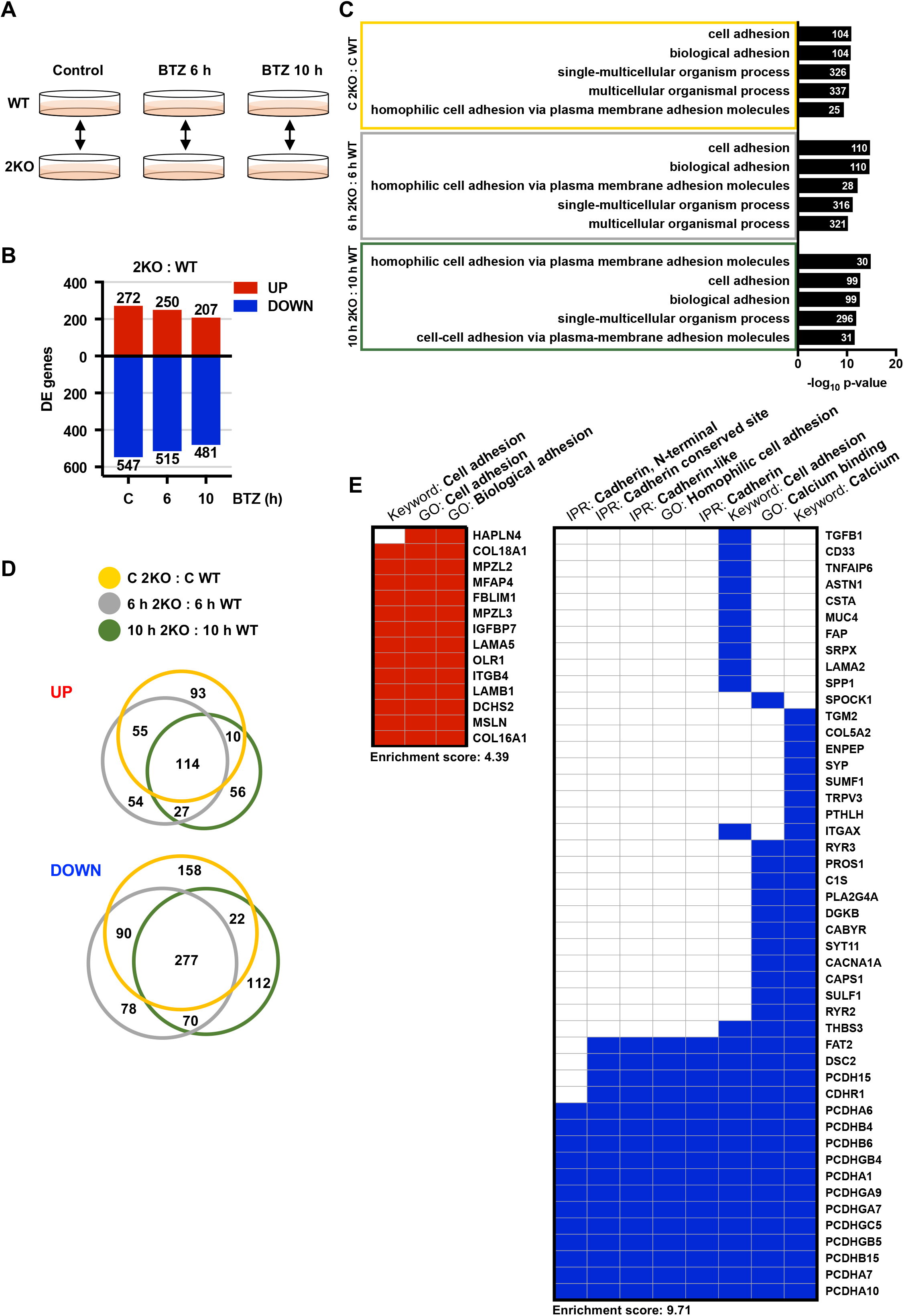
HSF2 impacts the expression of genes associated with cadherin-mediated cell-cell adhesion. **(A)** A schematic overview of the U2OS WT and HSF2 KO (2KO) comparison pairs. **(B)** Differentially expressed (DE) genes in 2KO : WT comparison pairs (control, 6 h, 10 h) were determined with Bioconductor R package Limma [48] (FC ≥ 3, FDR < 0.001). The upregulated and downregulated genes are indicated with red and blue bars, respectively. **(C)** Gene ontology (GO) terms were analyzed with topGO and GOstats packages in Bioconductor R. Biological processes from each comparison pair were ranked according to their p-values and the five most significantly changed GO terms are shown. The number of genes associated with a given term is indicated. **(D)** Venn diagrams presenting the interrelationship of significantly (FC ≥ 3, FDR < 0.001) upregulated or downregulated genes in 2KO : WT comparison pairs at control (orange), 6 h (grey), and 10 h (green) time points. Diagrams were generated using the BioVenn web application. **(E)** Gene term heatmap generated with DAVID Functional Annotation Clustering Tool based on the 114 upregulated (left panel) and the 277 downregulated (right panel) genes in 2KO cells in all treatment conditions as shown in panel D. Red and blue squares denote positive association between the gene and the keyword, Gene ontology (GO) term, or InterPro (IPR) term. Cluster enrichment score for upregulated gene cluster: 4.39; for downregulated gene cluster: 9.71.

To identify the adhesion molecules that are abnormally expressed in 2KO cells in both control and stress conditions, the gene set overlaps were examined with Venn diagrams. Among the comparison pairs, a total of 114 and 277 genes were upregulated and downregulated, respectively (Fig. 3D). Functional cluster annotation of the 114 upregulated genes with the DAVID analysis tool [50] confirmed the strong association to cell adhesion and extracellular matrix, including collagens (*COL16A1, COL18A1*) and laminins (*LAMB1* and *LAMA5*) (Fig. 3E left panel and S3). Interestingly, the 277 downregulated genes included members from many cadherin sub-families, for example protocadherins (*PCDHA1* and *PCDHA7*), desmosomal cadherins (*DSC2*), and Fat-Dachsous cadherins (*FAT2*), suggesting that the cadherin-mediated cell adhesion is extensively misregulated in 2KO cells (Fig. 3E, right panel). Cadherins are transmembrane adhesion molecules that mediate Ca^2+^-dependent cell-cell adhesion *via* the conserved extracellular cadherin domains [51]. The human genome encodes 110 cadherin genes, which together form the cadherin superfamily consisting of distinct cadherin sub-families [51]. The most prominent changes were detected in protocadherins, as 13 distinct protocadherin genes were significantly downregulated in 2KO cells grown either under normal growth conditions or exposed to BTZ-induced stress (Fig. 3E).

### Cells lacking HSF2 display abnormal cadherin expression

Since the cadherin genes appeared as a novel HSF2-dependent gene group and showed significant misregulation in multiple sub-family members, we examined the expression profiles of all cadherin superfamily genes in 2KO cells. Normalized gene expression data were used to generate a heat map encompassing all the cadherin superfamily genes encoded by the human genome. By comparing the expression profiles of WT and 2KO cells in control and BTZ-induced stress conditions, we observed a clear downregulation of the entire cadherin superfamily, as at least one member from every sub-family was found downregulated in 2KO cells. These included classical cadherins (*CDH2* and *CDH6*), desmosomal cadherins (*DSC2* and *DSG2*), CDH23-PCDH15 cadherins (*CDH12*), Fat-Dachsous cadherins (*FAT2* and *FAT4*), Flamingo cadherins (*CELSR1*), and Calsyntenins (*CLSTN2*) (Fig. 4A, Table S1). The most striking downregulation was detected in clustered α-, β-, and γ-protocadherins [52], of which 92% were found abnormally expressed in 2KO cells (Fig. 4A). Based on these results, we propose that the members of the cadherin superfamily are the main adhesion molecules downregulated in HSF2-depleted U2OS cells.

**Figure 4.**
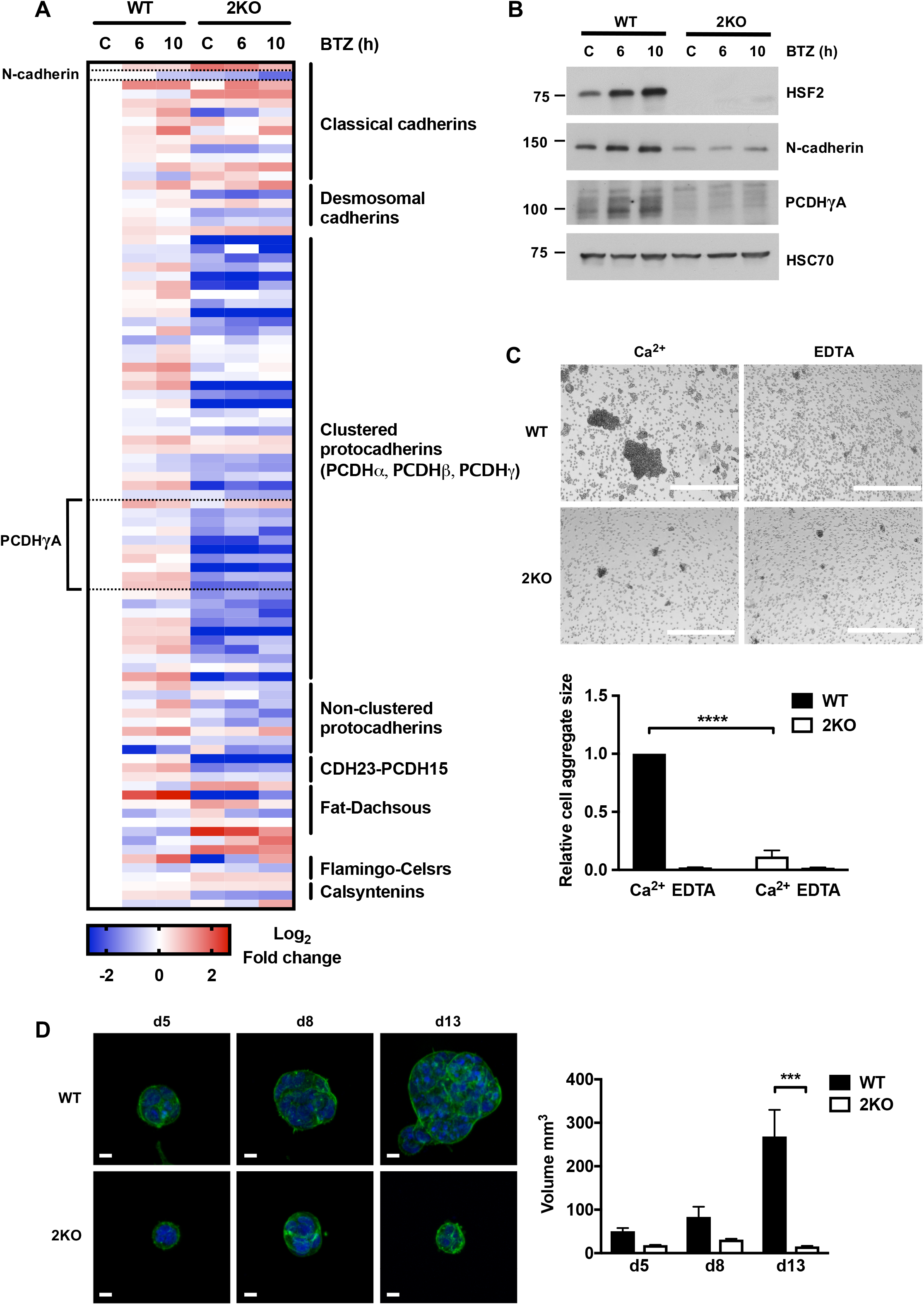
HSF2 controls cellular adhesion properties through cadherin superfamily proteins. **(A)** Normalized gene expression data from the RNA-seq analysis for cadherin superfamily genes, as defined in [51], was used to calculate the fold change of each gene in relation to respective expression in WT control sample. The data is presented as a heatmap of log2 transformed fold changes and it was generated with GraphPad Prism7. N-cadherin and protocadherin gamma subfamily A (PCDHγA) were chosen for further analyses. **(B)** Immunoblot analysis of HSF2, N-cadherin, and the members of PCDHγA. U2OS WT and 2KO cells were treated with 25 nM BTZ for 6 or 10 h. HSC70 was used as a loading control. **(C)** Cell aggregation assay of U2OS WT and 2KO cells suspended in cell aggregation buffer supplemented with 3 mM CaCl2 (Ca^2+^) or 3 mM EDTA. Cells were rotated for 2.5 h at 37°C and visualized with bright-field microscopy. Scale bar 1 mm. The size of the aggregates was quantified with ImageJ. The data is presented as mean values of three independent experiments + SEM. ****p < 0.0001. **(D)** Confocal microscopy images of U2OS WT and 2KO cells. Cells were cultured in 3D in Matrigel for 5, 8, and 13 days. At the indicated days, spheroids were fixed and F-actin was stained with Alexa 488-labeled phalloidin (green). DAPI was used to stain the nuclei (blue). Z-stacks of the spheroids were imaged with a spinning disc confocal microscope. The maximum intensity projection images represent the average spheroid size for each cell line at indicated time points from three biological repeats. Scale bar 10 μm. The volume of the spheroids was quantified with ImageJ with 3D Object Counter v2.0 plugin [65]. The data is presented as mean values of three independent experiments + SEM. ***p < 0.001.

For understanding the biological relevance of the RNA-seq analyses, we next determined the protein expression levels of N-cadherin (*CDH2*) and clustered γ-protocadherins (PCDHγA) by immunoblotting. As shown in Figure 4B, both N-cadherin and γ-protocadherins were found markedly downregulated also at the protein level in control and BTZ-treated 2KO cells (Fig. 4B). Since cadherins are essential in mediating Ca^2+^-dependent cell-cell contacts, we examined the functional impact of our observations with cell aggregation assay, where single cells are allowed to freely make cell-cell adhesion contacts in suspension. U2OS WT and 2KO cells were suspended in cell aggregation buffer supplemented with either CaCl2 or EDTA. WT cells supplemented with Ca^2+^ formed large cell aggregates, which were completely abolished in Ca^2+^-chelating conditions (EDTA) (Fig. 4C). In stark contrast, 2KO cells were unable to form cell aggregates even in the presence of Ca^2^+ (Fig. 4C), indicating that HSF2 is required to maintain Ca^2^+-dependent cell-cell contacts. Loss of distinct cell-cell adhesion molecules has been associated with cellular inability to form three-dimensional (3D) spheroids in ultra-low attachment round bottom plates [53]. We explored the spheroid forming capacity of U2OS WT and 2KO cells by culturing the cells in 3D extracellular matrix (ECM) using Matrigel. As expected, the spheroids originating from 2KO cells were significantly smaller than the WT spheroids (Fig. 4D), further confirming the functional impairment of cell-cell adhesion of these cells. A profound decline in the expression and function of cadherin superfamily proteins was observed also in 2KO#2 cells (Fig. S4), demonstrating that the alterations are not specific for a single cell clone. Altogether these results show, for the first time, that lack of HSF2 leads to disrupted cadherin expression at mRNA and protein levels, and results in functional deterioration of cadherin-mediated cell-cell adhesion.

### Impaired cell-cell adhesion sensitizes cells to proteotoxic stress

Albeit it is well-acknowledged that cadherins are essential mediators of tissue integrity and pivotal in regulating the development of multicellular organisms [51, 52], their impact on stress resistance has remained largely unexplored. To examine whether the observed impairment of cell-cell adhesion in 2KO cells can also contribute to the susceptibility of the cells to BTZ-induced stress, we aimed at restoring the cellular adhesion properties by re-introducing N-cadherin to 2KO cells. For this purpose, WT and 2KO cells were transfected with either Mock or N-cadherin plasmids and the N-cadherin expression was examined with immunoblotting (Fig. 5A). As shown in Figure 5A, we were able to restore the N-cadherin levels in 2KO cells. Moreover, re-introduction of N-cadherin resulted in a functional rescue of cell-cell adhesion in 2KO cells (Fig. 5B). Importantly, when exposed to BTZ, the 2KO cells expressing exogenous N-cadherin displayed significantly less cleaved PARP-1 than the Mock-transfected cells (Fig. 5C), suggesting that restoration of cell-cell adhesion can suppress cell death upon BTZ-induced proteotoxic stress.

**Figure 5.**
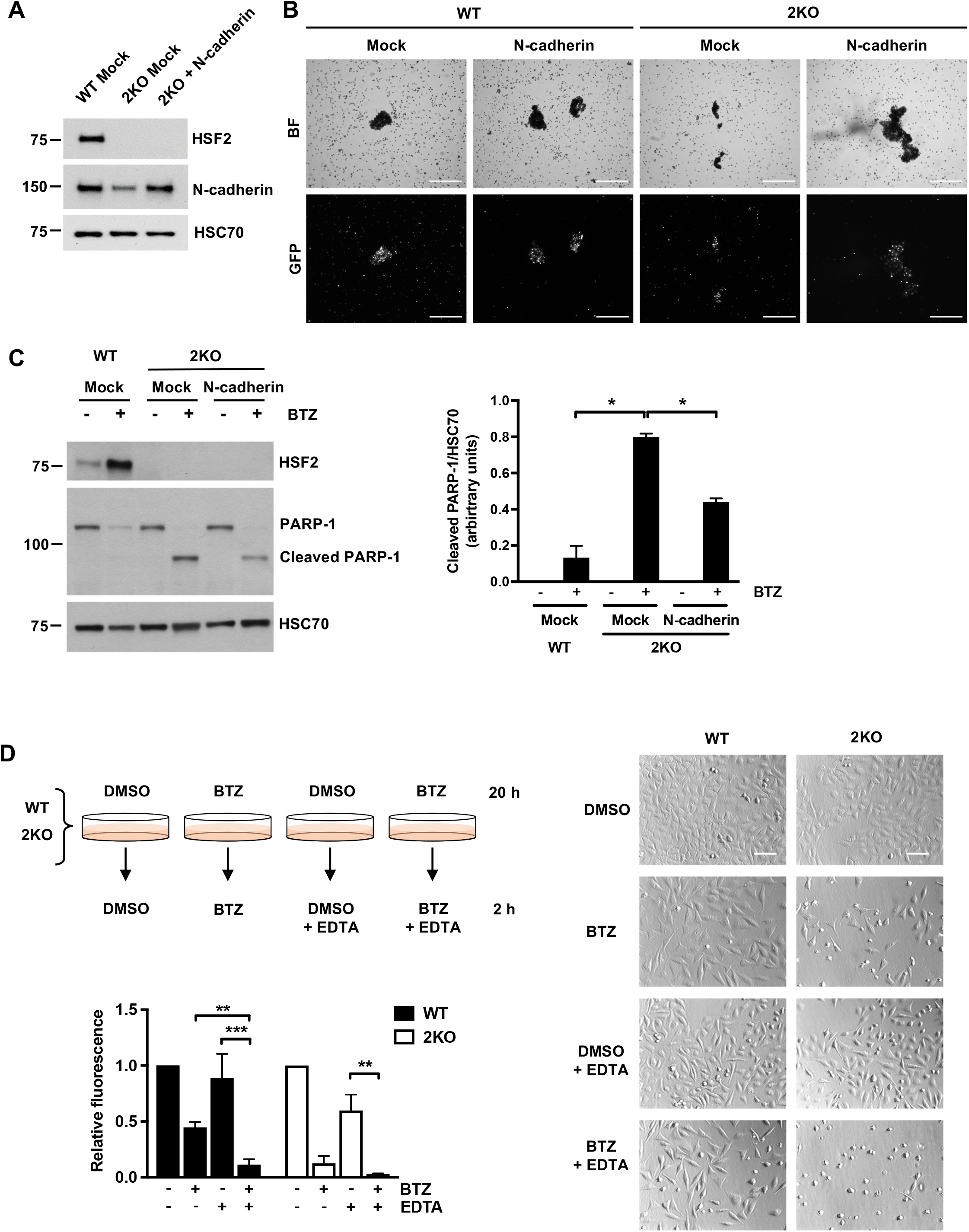
Impaired cell-cell contacts sensitize U2OS cells to BTZ-induced proteotoxic stress. N-cadherin levels in U2OS 2KO cells were restored to those in WT cells by transiently transfecting the cells with either Mock or N-cadherin plasmids co-expressing GFP (A-C), for details see Materials and Methods. **(A)** Immunoblot analysis of HSF2 and N-cadherin. HSC70 was used as a loading control. **(B)** Cell aggregation assay was performed as in Figure 4C. Cell aggregates were imaged with bright-field (BF) and fluorescence filters (GFP). Scale bar 500 μm. **(C)** For immunoblot analysis of HSF2 and PARP-1, cells were treated with 25 nM BTZ for 22 h. Control cells were treated with DMSO. HSC70 was used as a loading control. The amount of cleaved PARP-1 relative to respective HSC70 level was quantified with ImageJ. The data is presented as mean values of three independent experiments + SEM, *p < 0.05. **(D)** A schematic overview of the calcium-depletion experiments. U2OS WT and 2KO cells were first treated with or without 25 nM BTZ for 20 h in serum free growth medium after which the extracellular calcium was depleted with 3 mM EDTA and BTZ treatment continued for 2 h. Control cells were treated with DMSO. Cell viability was assessed with Calcein AM assay. Relative fluorescence was calculated against each respective control value that was set to 1. The data is presented as mean values of three independent experiments + SEM, **p < 0.01, *** p < 0.001. Statistical significance was determined with unpaired t-test. Cells were imaged with a bright-field microscope. Scale bar 100 μm.

All cadherin superfamily proteins are characterized by extracellular cadherin repeat domains, which mediate homophilic adhesion contacts between adjacent cells [54]. Stabilization of the extracellular domains is regulated by Ca^2^+, which binds to the interdomain regions of consecutive cadherin repeats and rigidifies the ectodomain structure. To be able to comprehensively investigate the role of cadherins in the cellular resistance to proteotoxic stress, we first treated the WT and 2KO cells with BTZ for 20 h to induce proteotoxic stress, after which the whole cadherin-mediated cell-cell adhesion program was destabilized by depleting the extracellular Ca^2^+ with 3 mM EDTA (Fig. 5D). Serum-free culture conditions were used to enable full depletion of extracellular Ca^2^+. Intriguingly, we observed that Ca^2+^-depletion intensified cell death, which was evidenced by the significantly reduced WT cell survival after treatment with both BTZ and EDTA (Fig. 5D). In contrast, the viability of 2KO cells was not markedly affected by BTZ treatment combined with Ca^2^+-depletion when compared to the BTZ treatment alone (Fig. 5D). Altogether, these results show that cadherin-mediated cell-cell adhesion is a key determinant of cell survival upon BTZ treatment and that destabilization of cadherin contacts predisposes U2OS cells to prolonged proteotoxic stress.

## DISCUSSION

Maintenance of cellular protein homeostasis (proteostasis) is crucial for the viability of all cells and organisms [13]. The heat shock response is critical for promoting proteostasis and it is under strict control of the heat shock transcription factors, HSFs. Until now, the comprehensive role of HSF2 in the cellular response to sustained proteotoxicity has remained unknown. We hypothesized that HSF2 is required for protecting cells against progressive accumulation of protein damage. We used the clinically relevant proteasome inhibitor Bortezomib (BTZ, PS-341, VELCADE^®^) as our experimental tool to induce long-term proteotoxic stress. BTZ treatment has been previously shown to upregulate HSF2 at both mRNA and protein levels in blood-derived human primary cells and to induce HSF2 binding at specific gene loci [35]. Our data confirmed the earlier results by showing that BTZ treatment leads to a remarkable increase in HSF2 protein levels also in malignant human cells. Moreover, we demonstrate that the amount of nuclear HSF2 is highly increased in BTZ-treated cells, strongly suggesting that HSF2 specifically responds to proteasome inhibition. We found that HSF2 is not only activated by BTZ-induced proteotoxicity, but it is absolutely essential for cell survival under these conditions. Based on our results, together with the reduced viability of *Hsf2^-/-^* MEFs treated with MG132 or exposed to febrile-range thermal stress [37, 41], we conclude that HSF2 is required to protect cells against progressive accumulation of damaged proteins.

Elevated protein levels of HSF1 and its phosphorylation on serine 326 were recently shown to be a prerequisite for multiple myeloma cell survival upon BTZ treatment [36, 55]. Therefore, we explored whether HSF2 depletion sensitizes cells to BTZ through misregulated HSF1, specifically, or the heat shock response in general. Neither difference in HSF1 levels nor serine 326 phosphorylation was detected between WT and HSF2-depleted cells treated with proteasome inhibitors. Moreover, the classical heat shock response, as characterized by the global upregulation of molecular chaperones, HSP90 co-chaperones, and polyubiquitin genes, was not compromised in cells lacking HSF2. These results indicate that HSF2 can promote cell survival independently of HSF1.

Previously, HSF2 has been shown to have only a modest effect on gene expression [2, 13, 41]. To our surprise, despite the stringent cutoff criteria (FC ≥ 3, FDR 0.001), we found a great number of genes displaying altered expression profiles in cells lacking HSF2. Among the most prominently misregulated genes were those belonging to the cadherin superfamily. Cadherins are a large group of transmembrane adhesion molecules, which mediate Ca^2+^-dependent cell-cell adhesion and thereby function as essential mediators of tissue integrity [51]. Lack of HSF2 led to a profound downregulation of cadherins both at mRNA and protein levels as well as functional impairment of cadherin-mediated cell-cell adhesion contacts. Our finding of abnormal cadherin expression already under normal growth conditions, together with the earlier results on HSF2 displaying DNA-binding capacity in the absence of stress [2, 15, 56], suggests that HSF2 has a physiological role in regulating cadherin gene expression. Excitingly, impaired migration and mispositioning of neurons have been shown to underlie the corticogenesis defects in *Hsf2^-/-^* mice [22, 23], and cadherin superfamily proteins are fundamental for correct neuronal migration [57]. Thus, it is tempting to speculate that the HSF2-dependent disruption of cadherin-mediated cell-cell contacts contributes to the abnormal corticogenesis of *Hsf2^-/-^* mice.

The downregulation of cadherin gene expression raises important questions about the mechanisms by which HSF2 regulates these genes. Genome-wide mapping of HSF2 binding sites has been previously determined with ChIP-seq in human K562 erythroleukemia cells [2] and in mouse testis [58]. Remarkably, both studies identified HSF2 occupancy on multiple cadherin superfamily genes. Since non-adherent K562 cells are deficient of endogenously expressed classical cadherins and distinct protocadherins [59], it is not surprising that HSF2 was found to occupy only the *CLSTN* gene under control growth conditions [2]. However, upon acute heat stress, HSF2 binding was observed at classical cadherins (*CDH4*), desmogleins (*DSG2*), Fat-Dachous cadherins (*DCHS2*), Flamingo cadherins (*CELSR2*), and CDH23-PCDH15 cadherins (*CDH23*) [2], demonstrating that multiple genes belonging to the cadherin superfamily can be targeted by HSF2 in human cells. In mouse testis, HSF2 was also shown to occupy several cadherin genes, including *CDH15, CDH5, CDH18, CDH13, FAT1, PCDH9, PCDH17,* and *PCDHA1* [58]. Altogether these results suggest that HSF2 is a key regulator of cadherin genes. Future studies are warranted for comprehensive analyses of direct HSF2 targets in cells and tissues across a variety of model systems.

Failure in maintenance of proteostasis is a hallmark of aging and neurodegenerative diseases [60]. Intriguingly, in a mouse model of Huntington’s disease, lack of HSF2 was shown to predispose mouse brain to poly-Q aggregates and reduce life span [41], suggesting that HSF2 is required to protect neurons from progressive accumulation of damaged proteins. Cell survival upon proteotoxic stress has been conventionally considered to depend on inducible transcriptional programs, such as the heat shock response or the unfolded protein response [1, 61]. However, in this study, we show that HSF2-dependent maintenance of cell-cell adhesion is an essential and novel determinant of proteotoxic stress resistance. Our results indicate that misregulation of distinct cellular properties already under normal growth conditions can sensitize cells to proteotoxicity. HSF1 and HSF2 might represent the two arms of the cellular resistance towards proteotoxic stress; HSF1 as an acute responder to protein damage and HSF2 as a factor maintaining the long-term stress resistance. Notably, a meta-analysis of transcriptional changes associated with Alzheimer’s disease and aging, HSF2 was identified as a gene commonly downregulated during aging [62]. Thus, it is possible that the age-associated downregulation of HSF2 and subsequent disruption of cadherin-mediated cell-cell adhesion, participates in sensitizing cells, such as neurons, to aggregate mismanagement.

## MATERIALS AND METHODS

### CRISPR-Cas9

Guide RNAs (gRNA) targeting the exon 1 of *HSF2* were designed using CRISPOR software (http://crispor.tefor.net/) and cloned into pMLM3636 gRNA expression plasmid (a gift from Keith Joung, Addgene plasmid #43860). U2OS cells were transfected with Cas9 and gRNA expression plasmids using Amaxa electroporation as recommended by the manufacturer (Lonza). The hCas9 plasmid was a gift from George Church (Addgene plasmid #41815). One week after transfections, cells were seeded at single cell density. Clones were genotyped by DNA sequencing of PCR products spanning the targeted region of the *HSF2* gene. The selected U2OS clones presented 3 different outframe mutations on *HSF2* exon 1, each corresponding to a different allele (Table S2). Guide RNA sequence targeting the 1^st^ AUG of the *HSF2* exon 1: 5’-UGCGCCGCGUUAACAAUGAA-3’. Following primers were used for PCR for validation: forward (hHSF2_Cr_ATG_F): 5’-AGTCGGCTCCTGGGATTG-3’ and reverse (hHSF2_Cr_ATG_R): 5’-AGTGAGGAGGCGGTTATTCAG-3’. For the experiments, we utilized HSF2 knock-out cell clone 1 (hereafter 2KO) and HSF2 knock-out cell clone 2 (hereafter 2KO#2).

### Cell culture and treatments

Cells were cultured in DMEM (Dulbecco’s Modified Eagle’s media, D6171, Sigma-Aldrich), supplemented with 10 % fetal calf serum, 2 mM L-glutamine and 100 μg/ml penicillin-streptomycin, and grown in 5 % CO2 at 37 °C. Proteasome inhibition was induced with Bortezomib (BTZ, sc-217785, Santa Cruz Biotechnology) or MG132 (Z-Leu-Leu-Leu-H, 317-V, Peptide Institute Inc.). Both inhibitors were diluted in DMSO (dimethyl sulfoxide, D8418, Sigma-Aldrich) and applied to cells in final concentrations indicated in the figures. Control cells were treated with DMSO only.

To induce protein misfolding with amino acid analogues, cells were starved for 17 h in L-arginine free culture medium (A14431-01, Gibco) supplemented with 10 % fetal calf serum, 2 mM L-glutamine and 100 μg/ml penicillin-streptomycin. Following that, L-Canavanine sulfate salt (C9758, Sigma) was applied to the cells in final concentrations indicated in the figure. Cells were treated for 3 or 6 h. After the treatments, cells were visualized with Leica phase contrast microscope, an EVOS FL Cell Imaging System (Thermo Fisher Scientific), or an Axio Vert A1-FL LED microscope (Carl Zeiss) and harvested for further analyses.

### Transfections

For transfections, 6 x 10^6^ U2OS WT or 2KO cells were suspended in 400 μl of Opti-MEM (11058-21, Gibco) and subjected to electroporation (230 V, 975 μF) in BTX electroporation cuvettes (45-0126, BTX). To downregulate HSF2 in WT cells HSF2 targeting shRNA and Scr vectors (previously described in [25]) were used. For restoring the protein levels of N-Cadherin in 2KO cells, a N-cadherin in pCCL-c-MNDU3c-PGK-EGFP vector co-expressing EGFP (a gift from Nora Heisterkamp, Addgene plasmid #38153) and pcDNA3.1/myc-His(-)A (used as Mock) were used. One day after transfection, cells were trypsinized, counted, re-plated, and let to recover for 24 h before BTZ treatments.

For cell aggregation assays, cells were transfected with GenJet (#SL100489-OS, SignaGen Laboratories) according to manufacturer’s instructions. Briefly, cells were plated 18 to 24 h prior to transfections to ensure 80 % confluency, and fresh culture media with supplements was added to the cells before transfections. The N-Cadherin encoding vector (described above) was used for transfections, and pEGFP-N1 (clonetech) was used as a Mock. The plasmids and the GenJet reagent were diluted in serum free media, and applied to the cells in a ratio of 1:2 (DNA:GenJet reagent). Cells were incubated with the DNA:GenJet mixture for 4 h, washed with PBS, and supplemented with complete culture media. Cells were let to recover for 24 h before the cell aggregation experiments.

### Immunoblotting

Cells were collected in culture media, washed with PBS (L0615, BioWest) and lysed in lysis buffer [50 mM HEPES, pH 7.4, 150 mM NaCl, 1 mM EDTA, 2 mM MgCl2, 1 % Triton X-100, 10 % glycerol, 1 x complete™ Protease Inhibitor Cocktail (04693159001, Roche Diagnostics), 50 mM NaF, 0.2 mM Na3VO4]. Protein concentration of the lysates was determined with Bradford assay. Equal amounts of cell lysates were resolved on 4-20 % or 7.5 % Mini-PROTEAN^®^ TGX precast gels (Bio-Rad) and the proteins were transferred to a nitrocellulose membrane. For HSF2 detection, membranes were boiled for 15 min in MQ-H_2_O and blocked in 3 % milk-PBS-Tween20 solution for 1 h at RT. Primary antibodies were diluted in 0.5 % BSA-PBS-0.02 % NaN3 and the membranes were incubated in respective primary antibodies overnight at 4 °C. The following antibodies were used: anti-GAPDH (ab9485, Abcam), anti-HSC70 (ADI-SPA-815, Enzo Life Sciences), anti-HSF1 (ADI-SPA-901, Enzo Life Sciences), anti-HSF1 pS326 (ab76076, Abcam), anti-HSF2 (HPA031455, Sigma-Aldrich), anti-HSP70 (ADI-SPA-810, Enzo Life Sciences), N-cadherin (04-1126, Millipore or ab76057, Abcam), anti-PARP-1 (F-2, sc-8007, Santa Cruz Biotechnology), anti-PanPCDHγA (75-178, NeuroMab), and anti-β-Tubulin (T8328, Sigma-Aldrich). Secondary antibodies were HRP-conjugated and purchased from Promega, GE Healthcare or Abcam. All immunoblotting experiments were performed at least three times.

### Immunofluorescence

2 x 10^5^ U2OS WT cells were plated on coverslips or MatTek plates (P35GC-.5-14-C, MatTek Corporation) 24 h before treatments. Cells were fixed with 4 % paraformaldehyde (PFA) for 15 min, permeabilized in 0.1 % Triton-X-100 in PBS and washed three times with PBST (PBS-0.5 % Tween20). Cells were blocked with 10 % FBS in PBS for 1 h at room temperature and incubated overnight at 4 °C with a primary anti-HSF2 antibody (HPA031455, Sigma-Aldrich), which was diluted 1:20 in 10 % FBS-PBS. Secondary goat anti-rabbit Alexa Fluor 488 (R37116, Invitrogen) was diluted 1:500 in 10 % FBS-PBS and the cells were incubated for 1 h in RT. Cells were washed three times with PBST, incubated with 300 nM DAPI diluted in PBS or mounted in Mowiol-DABCO mounting medium, and imaged with a 3i CSU-W1 spinning disc confocal microscope (Intelligent Imaging Innovations).

### Calcein assays

5 x 10^3^ U2OS WT and HSF2 KO cells were cultured in clear bottom 96-well plate (6005181, Perking Elmer) in complete culture media. Cells were treated with indicated concentrations of Bortezomib or MG132 for 22 h. For calcium-depletion, cells were treated with or without 25 nM Bortezomib for 20 h in serum free media. The extracellular calcium was depleted with EDTA in calcium free media and the Bortezomib treatment was continued for 2 h. Control cells were treated with DMSO. After treatments cells were washed with PBS and incubated for 30 min at 37 °C with Calcein AM (4892-010-K, R&D Systems) diluted 1:1000 in PBS. Fluorescence intensity was measured with Hidex Sense microplate reader (HIDEX Corp) with excitation and emission wavelengths 485 nm and 535 nm, respectively. Respective blank values were subtracted from the sample values and the viability of untreated control samples (WT or 2KO) was set to value 1. All the measurements were repeated at least three times.

### Cell aggregation assays

After trypsinization, 5 x 10^5^ U2OS WT and 2KO cells were suspended in 2 ml of aggregation assay buffer (137 mM NaCl, 5.4 mM KCl, 0.63 mM, Na_2_HPO_4_, 5.5 mM glucose, and 10 mM HEPES, pH 7.4) supplemented with either 3 mM CaCl_2_ or 3 mM EDTA. Cells were rotated for 2.5 h in 150 rpm at 37 °C, after which the aggregates were imaged with the EVOS FL Cell Imaging System (Thermo Fisher Scientific) or with an Axio Vert A1-FL LED microscope (Carl Zeiss). Cell aggregation assays were performed in biological triplicates. The area of the three biggest aggregates in each sample was measured with ImageJ (U. S. National Institutes of Health, Bethesda, Maryland, USA) for quantification purposes. All cell aggregation experiments were repeated at least three times.

### RNA-sequencing

2 x 10^6^ U2OS WT and HSF2 KO cells were plated and cultured overnight. Following day, cells were treated with 25 nM BTZ for 6 or 10 h. Control cells were treated with DMSO. Cells were collected, and total RNA was purified with AllPrep DNA/RNA/miRNA Universal Kit (80224, Qiagen) according to manufacturer’s instructions. Genomic DNA from mRNA columns was digested with DNAse I. The RNA library was prepared according to Illumina TruSeq^®^ Stranded mRNA Sample Preparation Guide (part #15031047). Briefly, poly-A containing mRNA molecules were purified with poly-T oligo magnetic beads and fragmented with divalent cations under elevated temperatures. For first-strand cDNA synthesis, RNA fragments were copied using reverse transcriptase and random primers. Unique Illumina TrueSeq indexing adapters were ligated to each sample. The quality and concentration of cDNA samples were analyzed with Advanced Analytical Fragment Analyzer and Bioanalyzer 2100 (Agilent, Santa Clara, CA, USA) and Qubit^®^ Fluorometric Quantitation (Life Technologies). Samples were sequenced with Illumina HiSeq 3000 (Illumina). All the experimental steps after the RNA extraction were conducted in the Finnish Microarray and Sequencing Center, Turku, Finland. RNA-sequencing was performed from four independent sample series.

### Bioinformatic analysis

The quality of the raw sequencing reads was confirmed with FastQC version 0.20.1 and aligned against the hg38 human genome assembly using TopHat2 version 2.1.0. Subreads version 1.5.0 was used to calculate gene level expression counts according to RefSeq-based gene annotations. The downstream analysis was carried out with R and Bioconductor. The data was normalized with TMM normalization method on the edgeR package. In all sample groups, the Spearman’s correlation value was above 0.97, indicating high reproducibility. Statistical testing between the sample groups was carried out using Bioconductor R package Limma [48] and the differentially expressed genes were filtered using fold change ≥ 3 and false discovery rate (FDR) of 0.001 as cutoff. Enrichment analysis for the differentially expressed (DE) filtered genes was performed with topGO and GOstats packages. GO terms in each comparison pair were ranked according to their significance (lowest p-value) and the most significantly changed terms were selected for the figures. Additional information regarding the term IDs can be found from http://www.geneontology.org.

### Quantitative RT-PCR (qRT-PCR)

RNA was isolated using a RNeasy mini kit (74106, Qiagen) according to the manufacturer’s instructions and quantified using a NanoDrop ND-1000 spectrophotometer (Thermo Fisher Scientific). Following that, 1 μg of total RNA was reverse transcribed with an iScript™ cDNA Synthesis Kit (#1708891, Bio-Rad). SensiFAST™ Probe Lo-ROX and SensiFAST™ SYBR^®^ Lo-ROX kits (Bioline) were used for qRT-PCRs that were performed with QuantStudio 3 Real-Time PCR system (Applied Biosystems, Thermo Fisher Scientific). All primers and probes were purchased from Sigma Aldrich. The following forward (f) and reverse (r) primers, and probes (pr) were used: fRNA18S5, 5’-GCAATTATTCCCCATGAACG-3’; rRNA18S, 5’-GGGACTTAATCAACGCAAGC-3’; prRNA18S5, 5’-FAM-TTCCCAGTAAGTGCG GGTC-BHQ-3’; fDSC2; 5’-ATCCATTAGAGGACACACTCTGA-3’; rDSC2, 5’-GCCACCGATCCTCTTCCTTC-3’; fPCDHA6, 5’-TGACTGTTGAATGATGGCGGA-3’; rPCDHA6, 5’-TCGGGTACGGAGTAGTGGAG-3’; fPCDHA10, 5’-AGGCATCAGCCAGTTTCTCAA-3’; rPCDHA10, 5’-GAGAGCAGCAGACACTGGAC-3’. The mRNA expression levels were normalized against the respective 18S RNA (*RNA18S5*) expression in a given sample. All reactions were run in triplicate from samples derived from four biological replicates.

### 3D cell culture and immunofluorescence

For examining cell growth in 3D, cells were embedded in growth factor reduced Matrigel (#356231, Corning) and cultured in Angiogenesis μ-slides (#81501, Ibidi) as described previously [63]. Briefly, wells were filled with 10 μl of Matrigel:culture medium (1:1 ratio), which was polymerized at 37 °C for 60 min. WT, 2KO, or 2KO#2 cells were seeded on top of the gel at a density of 700 cells per well, let to attach at 37 °C for 2 h, and covered with 20 μl of Matrigel:culture medium (1:4 ratio). The upper layer of Matrigel:culture medium was polymerized at 37 °C overnight, and appropriate humidity was ensured by adding droplets of MQ-H2O between the wells. Culture medium was changed every second day, and cell growth was monitored by imaging the cultures with a Zeiss Axio Vert A1-FL LED microscope (Carl Zeiss).

For immunofluorescence, spheroids were washed with 40 μl of PBS and fixed with 25 μl of 4 % PFA for 20 min at room temperature, followed by three washes with 40 μl of PBS. Spheroids were stained with 25 μl of 0.7 % Triton X-100, 1:500 Alexa Fluor 488 Phalloidin (#A12379, #A22287, Thermo Fisher Scientific), 300 nM DAPI in PBS at room temperature for 1 h. The stained spheroids were stored in PBS at 4 °C until imaging.

The spheroids were imaged as z-stacks with a 3i CSU-W1 spinning disc confocal microscope (Intelligent Imaging Innovations) using the same settings between the repeats. Spheroid volume was calculated based on the phalloidin staining using ImageJ v1.51n [64] software with the 3D Object Counter v2.0 (Bolte and Cordelieres, 2006) plugin. The threshold for background and object voxels was manually adjusted for each image in order to capture the whole volume of each spheroid.

### Visualization of the data

Heatmaps were generated with GraphPad Prism 7 Software (GraphPad Prism Software, La Jolla California USA, http://www.graphpad.com). Venn diagrams were generated with BioVenn web application (http://www.cmbi.ru.nl/cdd/biovenn/). DAVID Bioinformatic Resources 6.7 (https://david-d.ncifcrf.gov/home.jsp) was used for functional annotation clustering.

### Statistical analysis

All statistical analyses were performed with GraphPad Prism 7 Software (GraphPad Prism Software, La Jolla California USA, http://www.graphpad.com). The statistical significance was analyzed with two-way ANOVA and Holm-Sidak’s post-hoc test unless indicated differently.

## ACKNOWLEDGEMENTS

We are grateful to Joshua Weiner (University of Iowa, Iowa, US) for helpful discussions and advice and for providing us with the anti-PanPCDHγA antibody. All members of the Sistonen laboratory and Véronique Dubreuil from the Mezger laboratory are thanked for their valuable comments and critical review of the manuscript. Imaging was performed at the Cell Imaging Core, Turku Centre for Biotechnology, University of Turku and Åbo Akademi University. The instruments used in this project belong to the infrastructure of Biocenter Finland. We especially thank Markku Saari and Jouko Sandholm from the Cell Imaging Core of Turku Centre for Biotechnology for technical assistance and advice. The Bioinformatics unit of Turku Centre for Biotechnology is acknowledged for their expert assistance with the RNA-sequencing data analysis. The Bioinformatics unit is supported by University of Turku, Åbo Akademi University, and Biocenter Finland.

## DATA AVAILABILITY

The original data are available at Gene Expression Omnibus (GEO) database under accession no. GSE115973.

---

**Figure Supplement 1.**
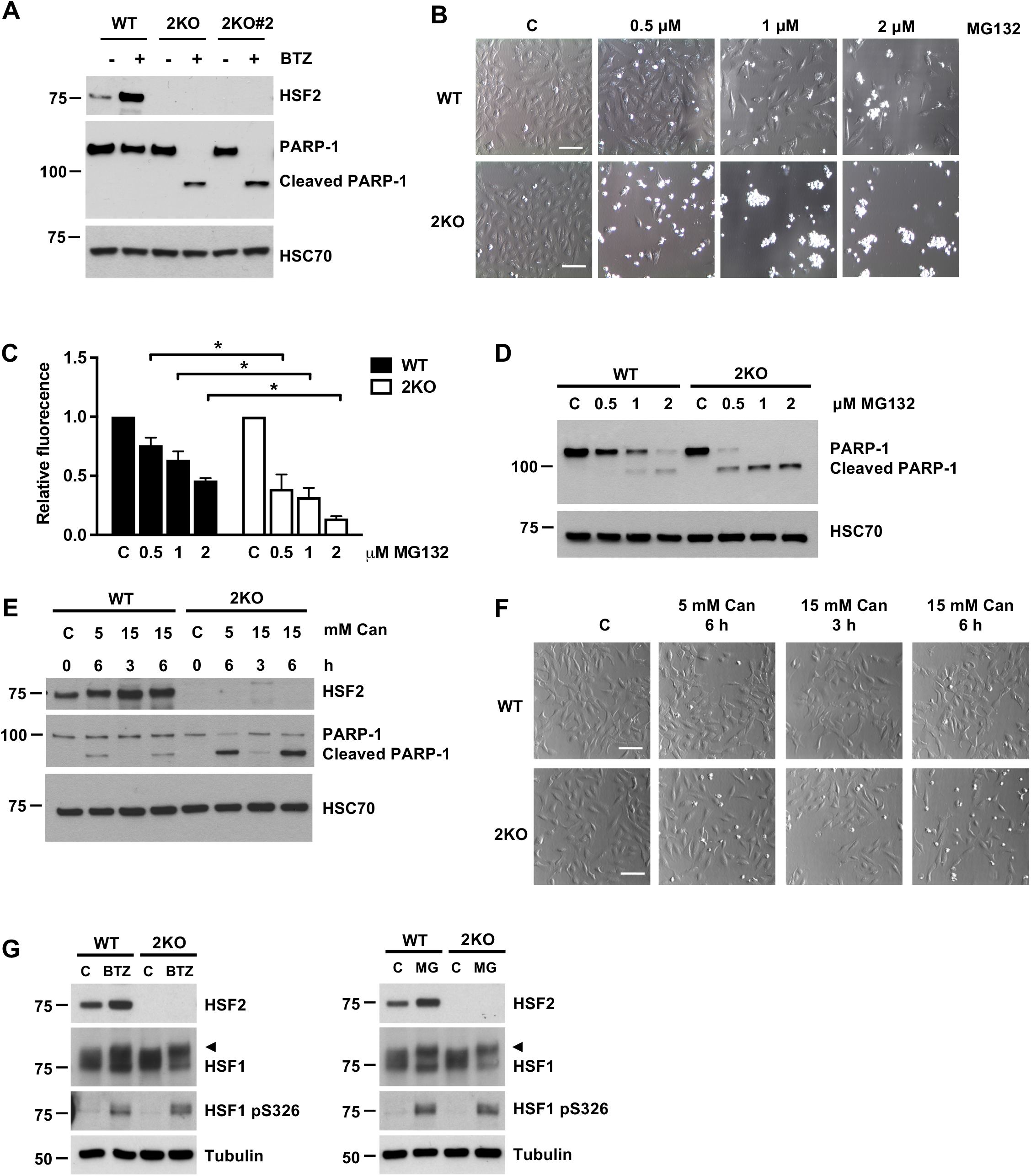
HSF2 depletion sensitizes cells to proteotoxic stress independently of HSF1 phosphorylation on serine 326. **(A)** Immunoblot analysis of HSF2 and PARP-1. U2OS WT, 2KO, and HSF2 knock-out clone 2 (2KO#2) cells were treated with 25 nM BTZ for 22 h. Control cells were treated with DMSO. HSC70 was used as a loading control. **(B)** Bright-field microscopy images of U2OS WT and HSF2 KO (2KO) cells treated with indicated concentrations of MG132 for 22 h. Control cells were treated with DMSO. Scale bar 100 μm. **(C)** Calcein AM assay of WT and 2KO cells treated as in A. Relative fluorescence was calculated against each respective control that was set to 1. The data is presented as mean values of at least three independent experiments + SEM, *p < 0.05. **(D)** Immunoblot analysis of PARP-1. Cells were treated as in A. HSC70 was used as a loading control. **(E)** Immunoblot analysis of HSF2 and PARP-1. U2OS WT and 2KO cells were starved in L-arginine free growth medium for 17 h after which the medium was supplemented with L-canavanine (Can) in indicated concentrations. Cells were treated in this growth medium for 3 or 6 h. HSC70 was used as a loading control. **(F)** Bright-field microscopy images of U2OS WT and 2KO cells treated as in E. Scale bar 100 μm **(G)** Immunoblot analysis of HSF2, HSF1 and HSF1 pS326. Cells were treated with either 50 nM BTZ or 10 μM MG132 (MG) for 6 h. Tubulin was used as a loading control. Arrowheads denote hyperphosphorylation of HSF1 [47].

**Figure Supplement 2.**
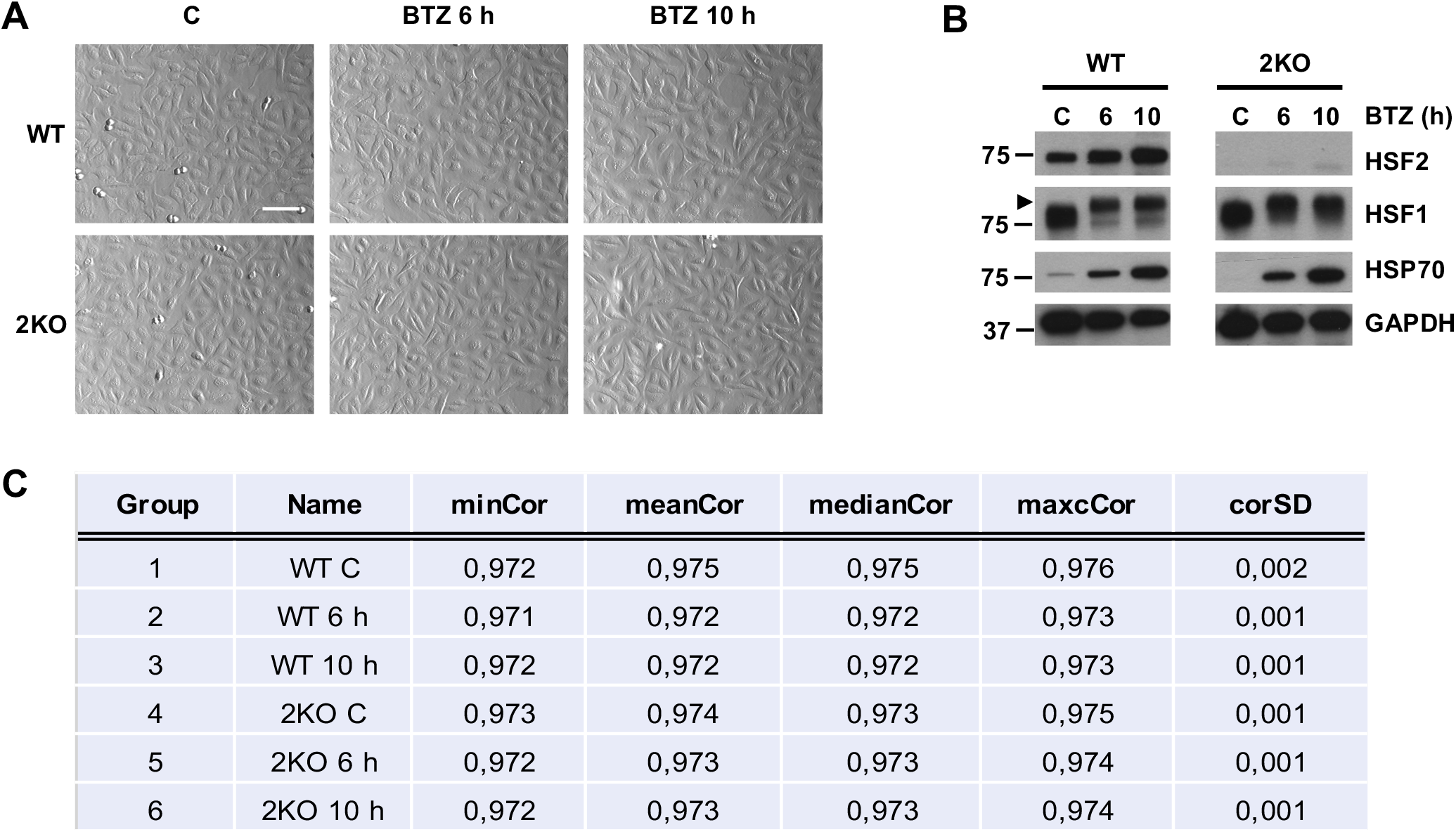
RNA-sequencing was performed in WT and HSF2-depleted U2OS cells. **(A)** Bright-field microscopy images of U2OS WT and HSF2 KO (2KO) cells treated with 25 nM BTZ for 6 or 10 h. Control cells were treated with DMSO. Scale bar 100 μm. **(B)** Immunoblot analysis of HSF2, HSF1 and HSP70. GAPDH was used as a loading control. Arrowheads denote hyperphosphorylation of HSF1 [47]. **(C)** Spearman’s correlation of RNA-seq sample replicates (n = 4).

**Figure Supplement 3.**
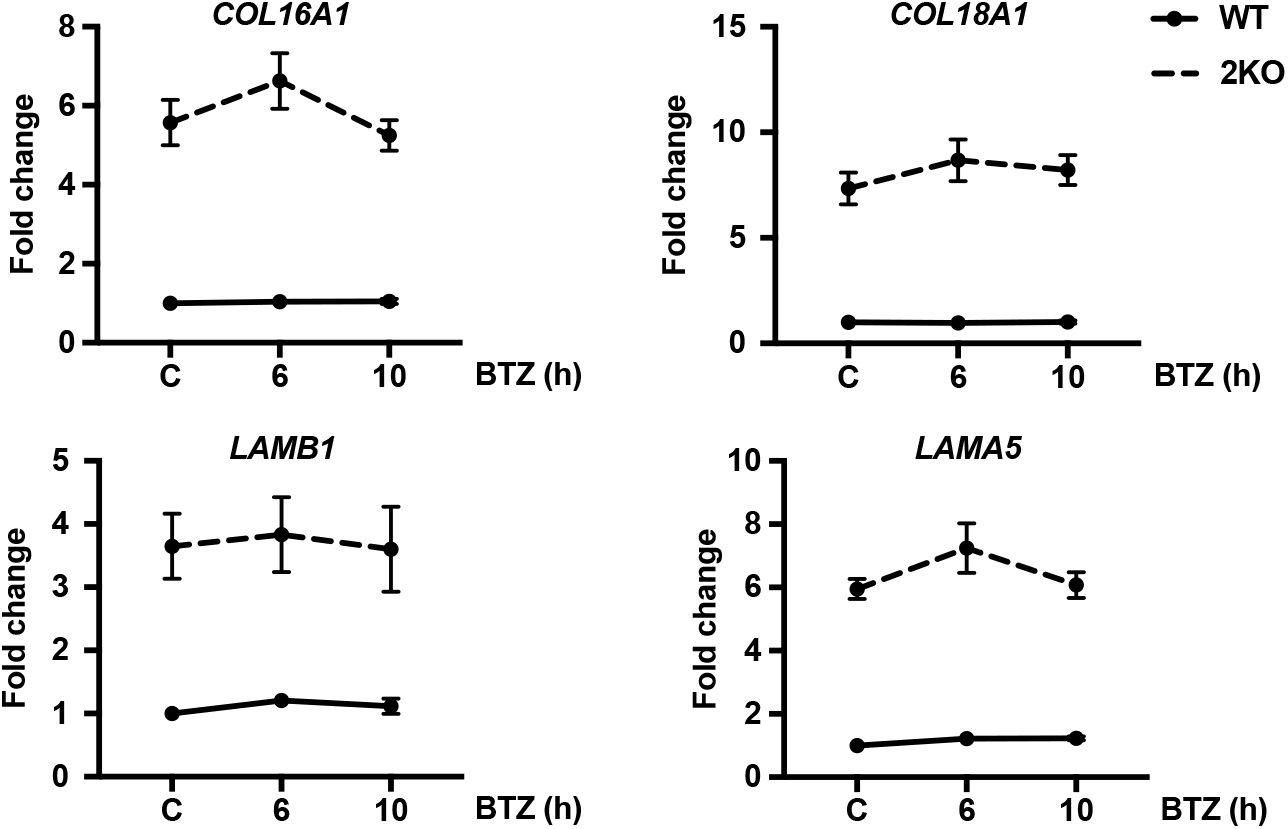
HSF2-dependent genes are linked to cell adhesion. mRNA expression levels of *COL16A1, COL18A1, LAMB1*, and *LAMA5* in control **(C)** and BTZ-treated WT and 2KO cells determined with RNA-seq. The data is presented as mean values ± SEM relative to WT control sample that was set to 1.

**Figure Supplement 4.**
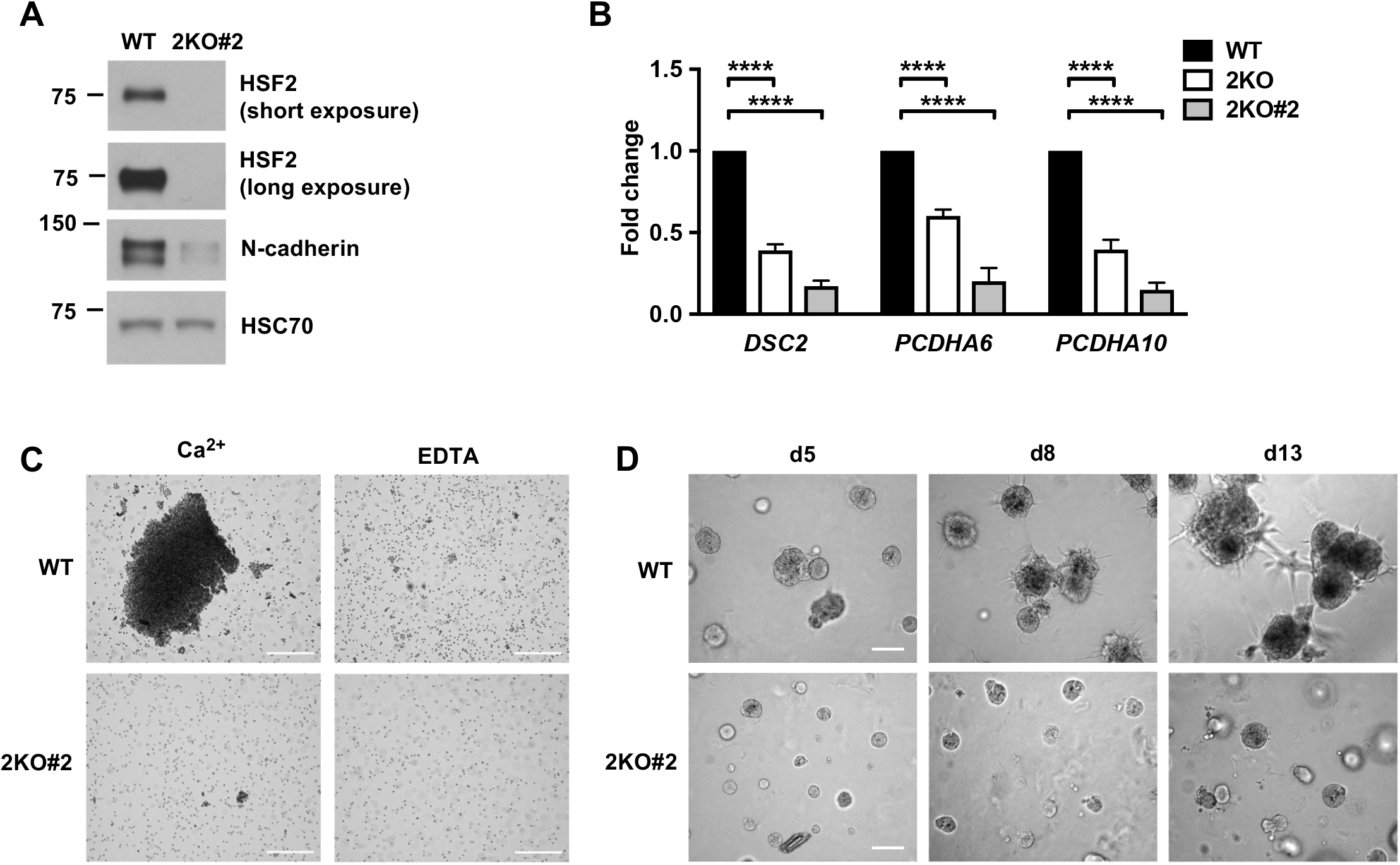
Cadherin misregulation and sensitization to proteotoxicity are not HSF2 knock-out cell clone specific. **(A)** Immunoblot analysis of HSF2 and N-cadherin in WT and 2KO#2 U2OS cells. HSC70 was used as a loading control. **(B)** mRNA expression of desmocollin 2 (DSC2), protocadherin α6 (*PCDHA6*), and protocadherin α10 (*PCDHA10*) of cadherin superfamily in WT, 2KO, and 2KO#2 cells. The mRNA levels were quantified with qRT-PCR. The data is presented as mean values of three independent experiments + SEM, ****p < 0.0001. **(C)** Cell aggregation assay of U2OS WT and 2KO#2 cells suspended in cell aggregation buffer supplemented with 3 mM CaCl2 (Ca^2^+) or 3 mM EDTA. Cells were rotated for 2.5 h at 37°C and visualized with bright-field microscopy. Scale bar 1 mm. **(D)** Bright-field microscopy images of U2OS WT and 2KO#2 cells cultured in 3D in Matrigel. The spheroids were imaged at days 5, 8, and 13. Scale bar 100 μm.

